# A brief report on the of mycobacterial proteins regulated by the loss of PtkA in Mtb-H37Ra

**DOI:** 10.1101/2023.09.10.556525

**Authors:** Swati Jaiswal, Kishore K Srivasatava

## Abstract

Kinase-phosphatase signaling plays an important role in cellular signaling pathways. The genome of *Mycobacterium tuberculosis*, (the causative agent of tuberculosis), is embossed with several kinases and phosphatases. Recently identified protein tyrosine kinase (PtkA) is a novel tyrosine kinase that phosphorylates protein tyrosine phosphatase A (PtpA) and regulates its secretion in the cytosol. PtkA is considered as a regulatory protein. In this study, the mycobacterial proteins regulated due to the *ptkA* gene deletion in Mtb-H37Ra have been studied and highlighted using two-dimensional (2D) gel electrophoresis coupled with MALDI TOF MS/MS analysis. We have identified 13 differentially expressed proteins that play an important role in mycobacterial intracellular survival and pathogenesis.

## Introduction

Tuberculosis is the deadliest disease after COVID-19 in the world (WHO 2022). It is caused by Mycobacterium tuberculosis (Mtb). To understand the biology of this slow-growing pathogen its complete genome was sequenced in 1998. The best-characterized strain of Mycobacterium tuberculosis, H37Rv, genome comprised of 4,411,529 base pairs, contains around 4,000 genes (Cole et al. 1998). Mycobacteria encode 11 serine threonine protein kinases (STPKs), 11 two component system and one tyrosine kinase (PtkA) which helps mycobacterium to evade host immune system and persist through the dormancy. The kinase and phosphatase pairs are best known to sense environmental signals and regulate intracellular survival during infection. The newly identified novel tyrosine kinase, PtkA (Bach et al. 2009), it lacks signature motif of Hank’s kinases and Walker A, A’ &Walker B motif which is known as the characteristic feature of prokaryotic tyrosine kinase, as reported by (Chao et al. 2014). PtkA activates by autophosphorylation at Y262. It is known to phosphorylates PtpA at two tyrosine residues Y128 and Y129 (Bach et al. 2009). The phosphorylations are necessary for the secretion of PtpA in the cytosol and regulates phosphatase activity of PtpA (Jaiswal S. et.al 2019). Here in this study author have investigated the mycobacterial proteins that are differentially regulated due to loss of ptkA using two-dimensional gel electrophoresis (2DE).

## Materials and Method

### Generation of ptkA knock out Mtb-H37Ra

The ptkA gene deletion mutant was created using the protocol mentioned here (Jaiswal S and Srivastava KK 2018).

### Preparation of Mtb lysate

Log phase cultures of wild type and *ptk A* knock out mycobacteria were harvested by centrifugation at 3000g for 10 min. The pellet was washed twice with PBS and resuspended in lysis buffer (Tris 20.0mM, NaCl 50.0mM, 1.0mM PMSF and protease inhibitor cocktail). The cells were then lysed by sonication at 25mm amplitude (10sec on, 10sec off) for 10 minutes per 10ml of culture used. The lysates were centrifuged at 12000g for 20 minutes to get a clear supernatant.

### 2D gel electrophoretic analysis

2DE was performed by standard procedures as described by manufacturer’s protocol (GE) (Jaiswal S and Srivastava KK 2018). Approximately 175.0μg of total protein was used for rehydrating (IPG strips pI 3-10 linear and 11cm length) (GE) by passive rehydration for 16h at 20°C. The rehydrated strips were run on Ettan IPGphor3 by gradually increasing the voltage across the IPG strips to at least 3500V and maintaining this voltage for several thousand Volt hours (25000-35000Vh). After focusing the proteins in the first dimension the 2nd dimensional SDS PAGE run. The strips were equilibrated in DTT containing equilibration buffer and then in the IAA containing buffer before being placed on 12.0% polyacrylamide gel for molecular weight dependent run in 1X Tris-Glycine-SDS buffer. The gel was run O/N at 16mA current (per gel) at 16°C. The gel was stained using Coomassie Brilliant Blue R-250. The destained gels were scanned using Image Scanner III (GE) and compared using IMP7 software (GE).

### MALDI TOF - MS (Matrix-assisted laser desorption/ionization-time of flight mass spectrometry) Analysis

The gel pieces were excised for in-gel tryptic digestion and destained using 100.0mM ammonium bicarbonate in 50.0% acetonitrile solution. Then were dehydrated using a solution containing 2:1 mixture of acetonitrile and 50.0mM ammonium bicarbonate for 5 minutes followed by a 2-minute wash with 25.0mM ammonium bicarbonate. This cycle was repeated 4 times and then the gel pieces were vacuum dried. The gel pieces were trypsin (0.02 μg/μL) digested at 37°C O/N. Later, the gel pieces were vortexed vigorously and the supernatant containing the peptides was mixed with α-Cyano-4-hydroxycinnamic acid (CHCA) matrix in a 1:1 ratio. The spotted peptides were analyzed using AB SCIEX 4800 MALADI TOF-MS analyzer (Applied Biosystems, USA) with 4000 series explorer v 3.5 software.

## Result

### Comparative analysis of differentially expressed proteins-

The whole cell lysates of wild type and ptkA deleted mutant Mtb was prepared as described in the methods section. The clear supernatant was focused based on their pI value on one dimensional gel strip. Further these proteins were separated through 2-DE shown in Figure 1. 25 spots were differentially expressed in ΔptkA Mtb in comparison to wildtype Mtb (Figure 1). Out of 25 spots, 19 spots were identified by MALDI-TOF MS analysis. The mycobacterial proteins regulated by deletion of ptkA gene in Mtb-H37Ra are listed here (Table 1).

**Table 1.**
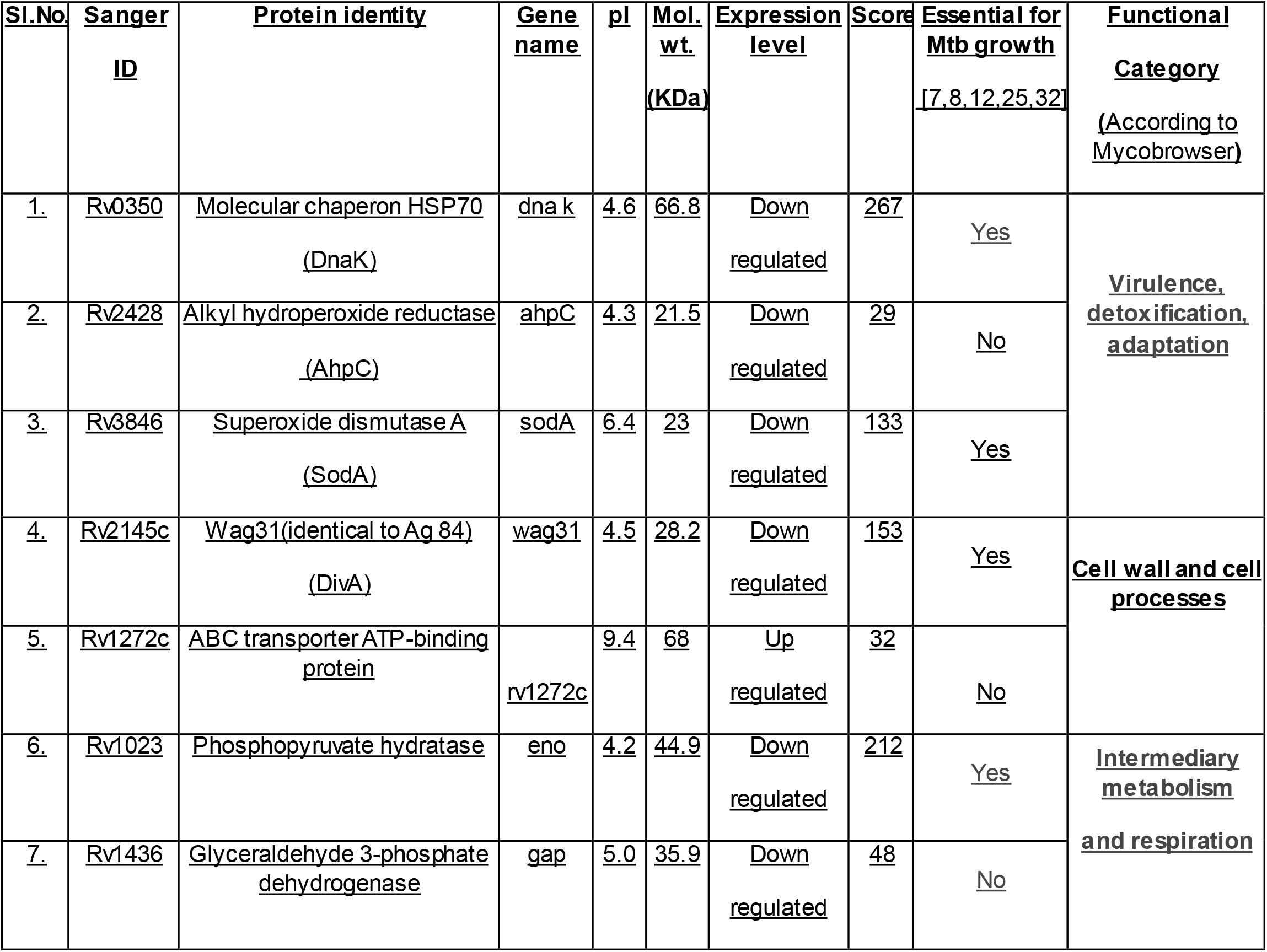
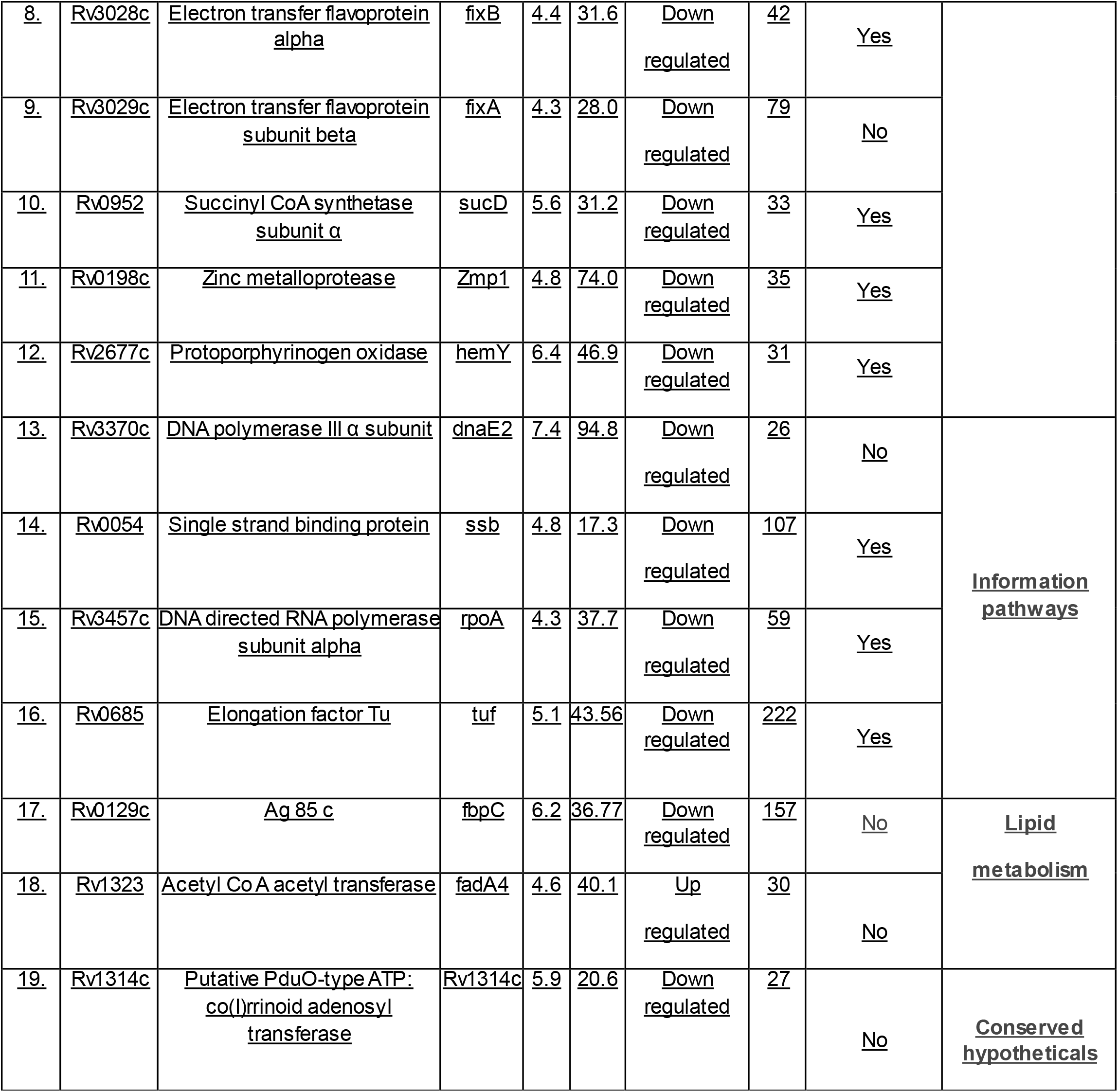
Mycobacterial Proteins regulated by loss of *ptkA* gene in Mtb-H37Ra.

**Figure 1.**
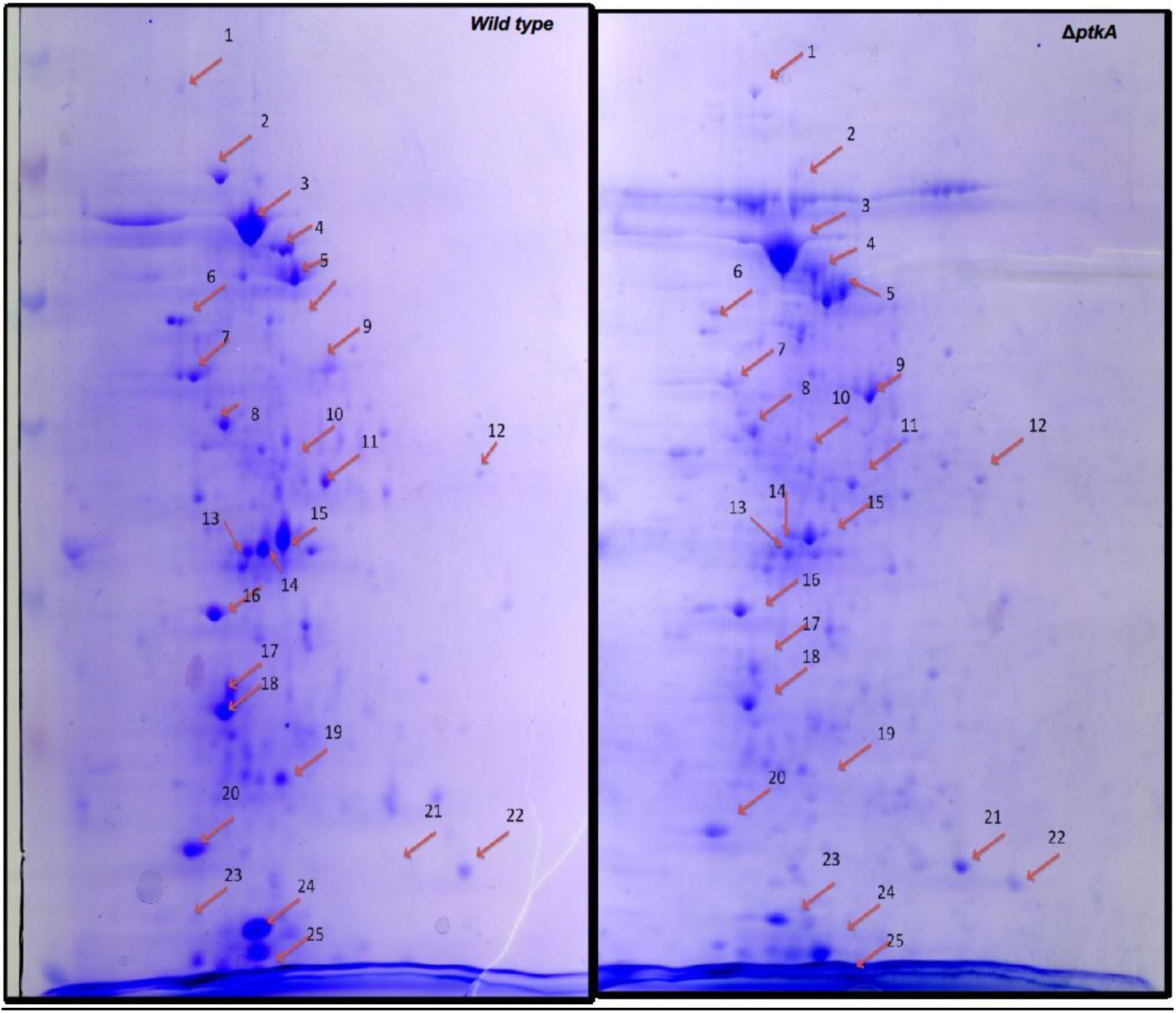
Two-dimensional gel electrophoresis of differentially expressed proteins in *ptk A* deleted mutant in comparison to wild type H37Ra.

### Proteomic profiling of the ΔptkA Mtb-

The proteomic analysis of *ptk A deleted mutant Mtb* have shown that 17 proteins are down regulated, and 2 proteins are upregulated in the ΔptkA Mtb. The proteins that have similar functions are grouped together and shown in Figure 2. The proteins that are downregulated in the ΔptkA Mtb are involved in relieving stress (superoxide dismutase and alkyl hydroperoxide reductase); metabolism (Phosphopyruvate hydratase, Glyceraldehyde 3-phosphate dehydrogenase, Electron transfer flavoprotein alpha and beta, Succinyl CoA synthetase subunit α, Ag 85 c, Protoporphyrinogen oxidase, Rv1314c and Zinc metalloprotease) maintaining cell processes (Wag31, DNA directed RNA polymerase subunit alpha, DNA polymerase III α subunit, Single strand binding protein, and Elongation factor Tu). Other two proteins that are upregulated are 1) involved in cellular transport; ABC transporter ATP-binding protein and 2) Acetyl Co A acetyl transferase, under studied in Mtb, known to involved in lipid metabolism.

**Figure 2.**
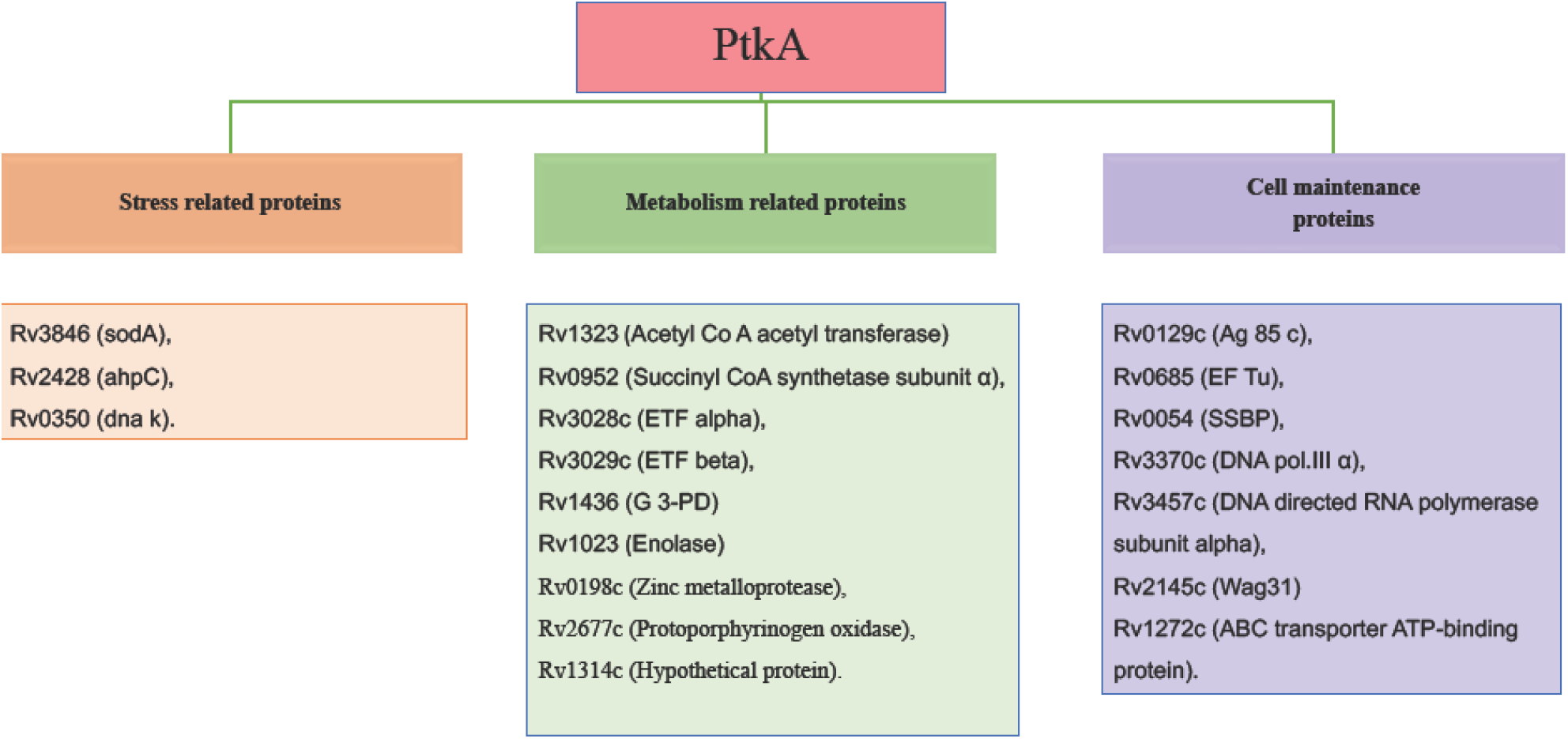
Mycobacterial proteins regulated by the loss of *ptk A* gene in Mtb-H37Ra are grouped in different categories based on their functions.

## Discussion

PtkA is a regulatory protein which modulates host proteins and help mycobacteria to survive inside the host. In this study we aim to investigate the proteins that are directly or indirectly regulated by ptkA. We used 2DE technique to identify the proteins that were differentially regulated (Gu Sheng et.al, 2003; Sharma D. et.al. 2015). Based on their functional characterization, PtkA seems to be involved in various functions such as in the stress conditions, affect ATPase activity, involved in the regulation of metabolic pathways (glycolysis & kreb’s cycle), maintain cell wall and cell processes, involved in oxidative stress response, and could also be involved in DNA recombination and repair. PtkA regulates protein biosynthesis and thought to be involved in active transport of drugs across the membrane.

### I. Stress related proteins-

PtkA deletion mutant is known to enhance secretion of **TrxB2 (thioredoxin reductase)** (Wong D. et al. 2019), a secretory protein and essential for Mtb growth. The mechanism is not known yet. Along with this it has found regulating other stress induced proteins-

**Superoxide dismutase A: sodA (Rv3846)**, is an essential protein for growth of Mtb-H37Rv (Griffin et al., 2011; DeJesus et al. 2017; Minato et al. 2019). It is present in the culture filtrate and plays a significant role in resisting oxidative stress (Liao D et.al, 2013).

**Alkyl hydroperoxide reductase; ahpC (Rv2428)** is identified in the membrane fraction of Mtb (Gu et al., 2003). In addition to several other proteins (secA2, mshA and cysH) these are implicated in ROS and RNS detoxification (Ehrt and Schnappinger 2011). These proteins were downregulated in the ptkA deleted Mtb-H37Ra.

### II. Metabolism related proteins-

**Acetyl Co A acetyl transferase, fadA4 (Rv1323)** was upregulated. It is not much studied yet but supposed to involve in lipid degradation (Mycobrowser database).

**Phosphopyruvate hydratase or Enolase (Rv1023)**, is an essential glycolytic enzyme that catalyzes the conversion of 2, phosphoglycerate (PGA) to phosphoenol pyruvate (PEP). Recently it has been reported to be involved in the formation of biofilm, however the mechanism is unknown (Kumar A et.al., 2023). Enolase has also been identified as an immunogen and a vaccine candidate for several pathogens including Mtb (Rahi A. et.al. 2017).

**Glyceraldehyde 3-phosphate dehydrogenase (GAPDH); gap (Rv1436)**, is present on the surface of Mtb. It has been reported to play an important function in iron acquisition from the host. Rv1436 binds with host transferrin and transport iron to the bacterium (Boradia V.M. et.al. 2014). Additionally, gap binds to lactoferrin with more affinity (Malhotra H. et.al. 2017).

**Electron transfer flavoprotein alpha; fix B (Rv3028c), and Electron transfer flavoprotein subunit beta; fix A (Rv3029c)** are essential genes for in vitro growth in H37Rv strain (Sassetti et al., 2003; DeJesus et al. 2017). These genes transfer the electrons to the main respiratory chain via ETF-ubiquinone oxidoreductase (ETF dehydrogenase).

**Succinyl CoA synthetase subunit α; SucD (Rv0952)**, is an essential protein for growth of Mtb-H37Rv (Griffin et al., 2011; DeJesus et al. 2017; Minato et al. 2019). It is identified in the cytosol, cell wall, and cell membrane fractions of Mtb (Mawuenyega et al., 2005). In the absence of oxygen, SucD is the only mitochondrial enzyme capable of ATP production via substrate level phosphorylation. Inorganic phosphate (iP), is a signaling molecule capable of activating oxidative phosphorylation, binds SucD in a non-covalent manner and enhances its enzymatic activity (Philips D. et.al, 2010).

**Ag 85 c (Rv0129c)** belongs to the Ag85 protein family (Ag A/ B/ C), and important to Mtb cell envelope biogenesis (Warrier T. et.al. 2012). The Ag85C transfers mycolic acid to the AG complex in vivo (Jackson M. et. al.1999). Inhibiting Ag85 complex is an attractive drug target. Cyclipostins and cyclophostin (CyC) analogs are shown to inhibit the antigen 85C both in vitro and in vivo (Viljoen A et.al 2018).

**Protoporphyrinogen oxidase; hemY (Rv2677c)**, is an essentialgene for in vitro growth of Mtb-H37Rv (Griffin et al., 2011; DeJesus et al. 2017; Minato et al. 2019) involved in heme and porphyrin biosynthesis, were found downregulated in the ptkA deleted Mtb-H37Ra.

**Zinc metalloprotease; Zmp1 (Rv0198c)**, a protein present in the culture filtrate (de Souza et al., 2011). Zmp1 is essential for the intracellular survival of the bacteria and possibly impairs inflammasome activation and phagosome maturation (Master et al., 2008; Johansen et al., 2011). The protein expression was downregulated in the ptkA deleted mutant. PtkA regulates expression of Zmp1, which is a Mtb specific immune stimulant and proposed as a potential vaccine candidate or a disease marker (Vemula MH et.al., 2016).

### III. Cell wall and cell processes-

**Wag31(or Ag 84); wag31 Rv2145c**, is an essential gene for invitro growth of Mtb (Minato et al. 2019; DeJesus et al. 2017; Sassetti et al., 2003; Griffin et al., 2011). It is known to modulate the host cytokine secretion of IL-10 and IL-17 and STAT3. (Park HS et.al. 2021; Samten B. et.al, 2016). It is a secretory protein and plays a crucial role in cell division and cell wall synthesis in mycobacteria (Meniche X et.al, 2014; Kang et al. 2005; Nguyen et al. 2007).

**DNA directed RNA polymerase subunit alpha; rpoA (Rv3457c)**, is an essential protein for invitro growth of Mtb (Minato et al. 2019; DeJesus et al. 2017; Sassetti et al., 2003; Griffin et al., 2011). It is identified in the culture filtrate, membrane protein fraction, and whole cell lysates of Mtb (de Souza et al., 2011). It catalyzes the transcription of DNA into RNA using the four ribonucleoside triphosphates as substrates and plays important role in the mycobacterium survival.

**DNA polymerase III α subunit; dnaE2 (Rv3370c)**, also known as error prone polymerase. The frequent use of antibiotic generates antibiotic persisters (AP) in the cultures. These AP Mtb experience oxidative stress and activates SOS response resulting in upregulation of DnaE2 and develop resistance to the lethal dose of ciprofloxacin or rifampicin (Salini S. et.al, 2022).

**Single strand binding protein; ssb (Rv0054)** is essential for mycobacterial intracellular survival (Minato et al. 2019; DeJesus et al. 2017). It is identified in the culture filtrate, membrane fraction and whole cell lysate of Mtb. This protein is essential for replication of the chromosome. It is also involved in DNA recombination and repair. MtbSSB binds with MtbDnaB (important in the initiation and the extension stage of DNA replication. MtbSSB assist in the loading of MtbDnaB on the DNA replication fork in Mtb (Zhang H et.al. 2014).

**Elongation factor Tu (Ef-Tu); tuf (Rv0685)** was found downregulated in the ptkA deleted Mtb-H37Ra. It is essential for in vitro growth (Minato et al. 2019; DeJesus et al. 2017; Griffin et al., 2011). It is identified in the cytosol, cell wall, and cell membrane fractions of Mtb-H37Rv. Ef-Tu is responsible for the selection and binding of the cognate aminoacyl-tRNA to the acceptor site on the ribosome. Its activity is dependent on its interaction with GTP. It has been reported that Protein kinase B (PknB), a STPK, phosphorylates MtbEf-Tu, including Thr118, which is required for optimal activity of the protein. The phosphorylation reduces its interaction with GTP (Sajid A. et.al. 2011).

Increased expression of the **ABC transporter ATP-binding protein (Rv1272c)** in the *ptk A* deletion mutant was observed. Braibant et al. in 2000 have reported that they have identified at least 37 complete and incomplete ABC transporters. They called them importers and exporters. These transporters play critical role in Mtb survival within the host. In comparison to the E. coli and B. subtilis, Mtb seems to have transporters that are important for virulence, cell attachment, drug transport, importing inorganic phosphate, necessary for survival inside the host (phagosomes, granulomas, caseum, etc.). Later, Rv1272c is shown to transport triacylglycerols (TAGs) when expressed in E. coli (Martin A et.al. 2018). Authors argues that it might be involved in the dormancy associated TAG synthesis in the Mtb.

The **conserved protein Rv1314c** is not essential for the growth (Sassetti CM et al. 2003; Griffin et.al. 2011), and localized in the culture filtrate, membrane protein fraction and whole cell lysates (de Souza et al., 2011). It is identified as putative PduO-type ATP: co(I)rrinoid adenosyl transferase as essential for B_12_ assimilation (Gopinath K et.al. 2013; Jinich A et.al. 2022). It interacts with other adenosyl transferase including Rv2850c and Rv1496, a transport system kinase, (probably GTPase) (STRING Version 11.5).

The proteins that are essential for mycobacterial growth are downregulated in the ptkA deletion mutant suggest that PtkA is an important regulatory protein for mycobacterial survival. The pattern of protein spots in complemented and wild type strains is similar (data not shown here).

PtkA interacts with several Mtb proteins and regulate their expression and thus plays important role in Mtb physiology. It phosphorylates and augment PtpA secretion, which is an important protein within macrophage after Mtb infection suggest that PtkA may have significant effect on host protein and regulate the survival of Mtb intracellularly.

## Acknowledgement

I am thankful to CSIR-Central Drug Research Institute for providing the core facility to carry out my research. The work has been funded through CSIR network projects Splendid (BSC0104) and UNDO (BSC0103).

## Author’s Contribution

SJ has performed the experiment, analyzed the data and written the manuscript. KKS has conceptualized the work, reviewed and finalized the manuscript.

## Conflict of Interest

The authors declare no conflict of interest

## Acknowledgements

We thank Director, CSIR-CDRI for the support. The CSIR-CDRI communication number allotted to this manuscript is (to be included after acceptance). The work has been supported through CSIR-CDRI budget heads MLP0107 and GAP0120. SJ is the recipient of CSIR JRF.

## Notes

### Competing Interest Statement

The authors have declared no competing interest.

